## Supplementary figures and images for "The domesticated transposon protein L1TD1 associates with its ancestor L1 ORF1p to promote LINE-1 retrotransposition"

### Supplemental Figure S1

#### Supplementary Figure S1

A

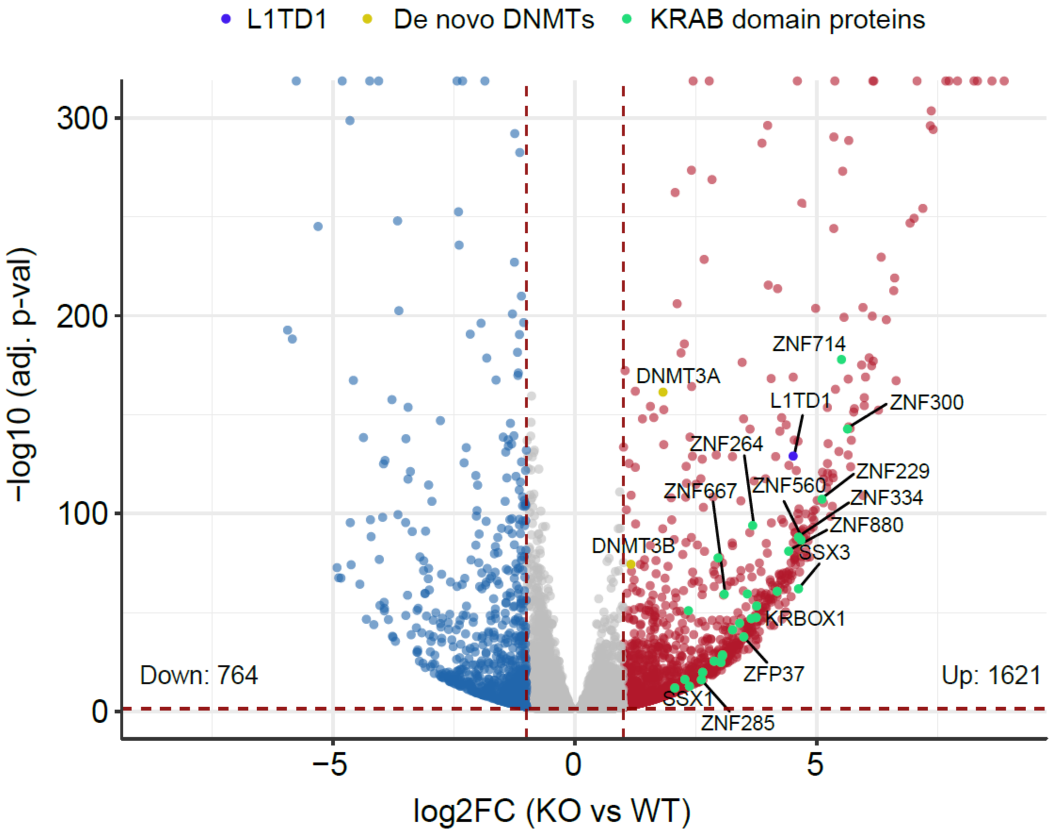


B

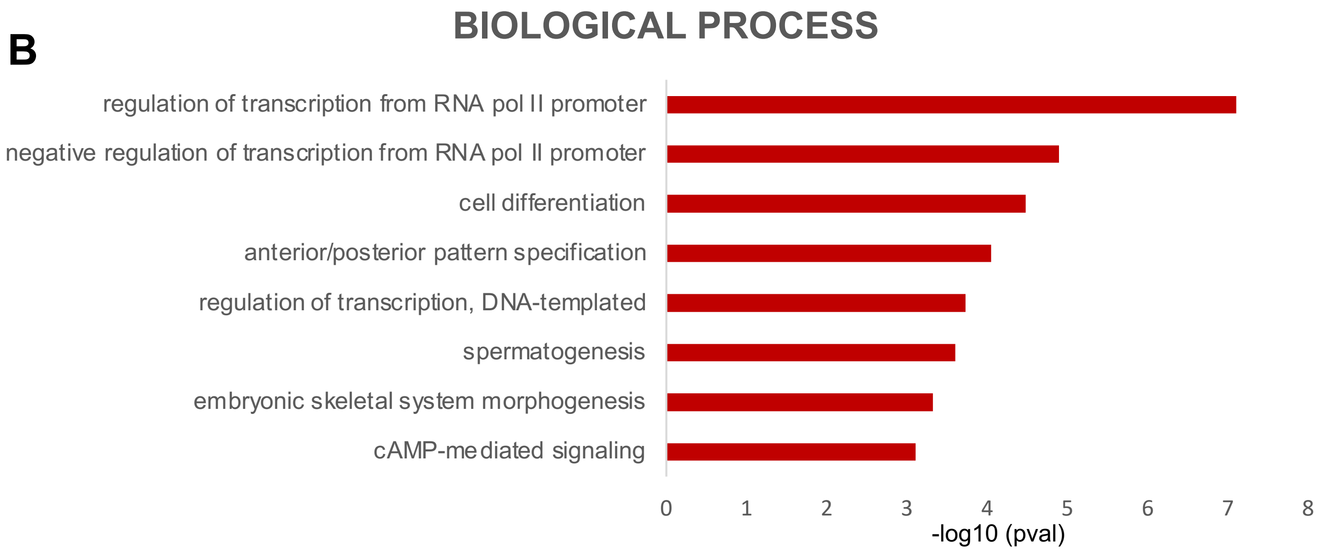


C

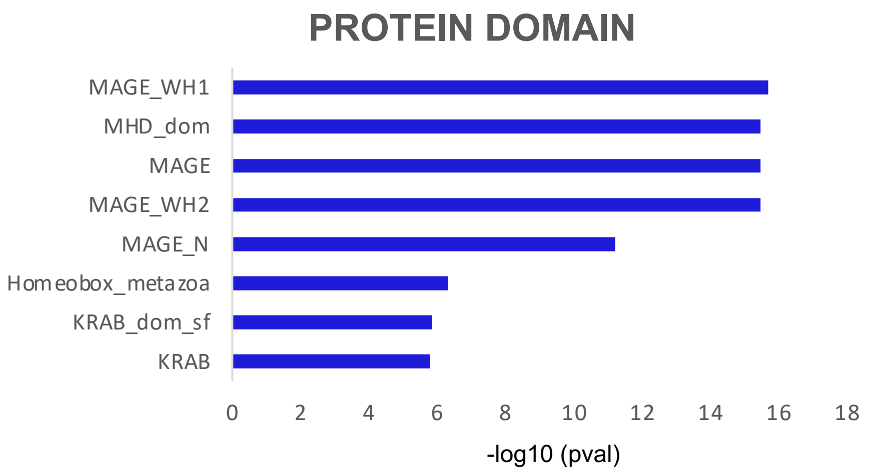

### Supplemental Figure S2

##### Supplementary Figure S2

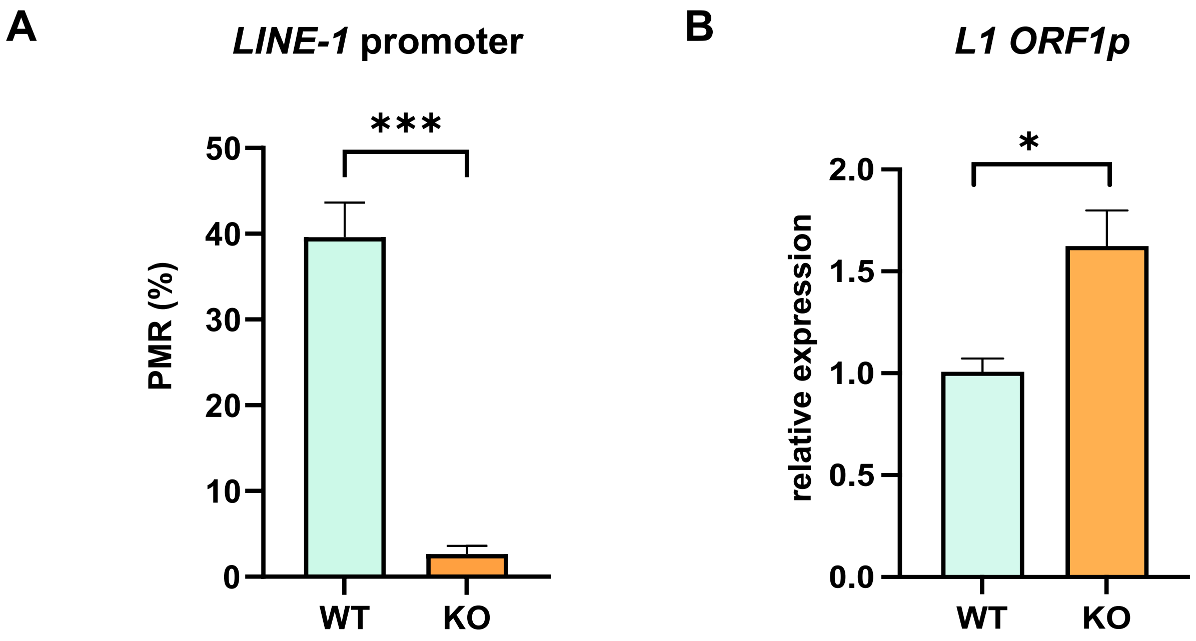

### Supplemental Figure S3

#### Supplementary Figure S3

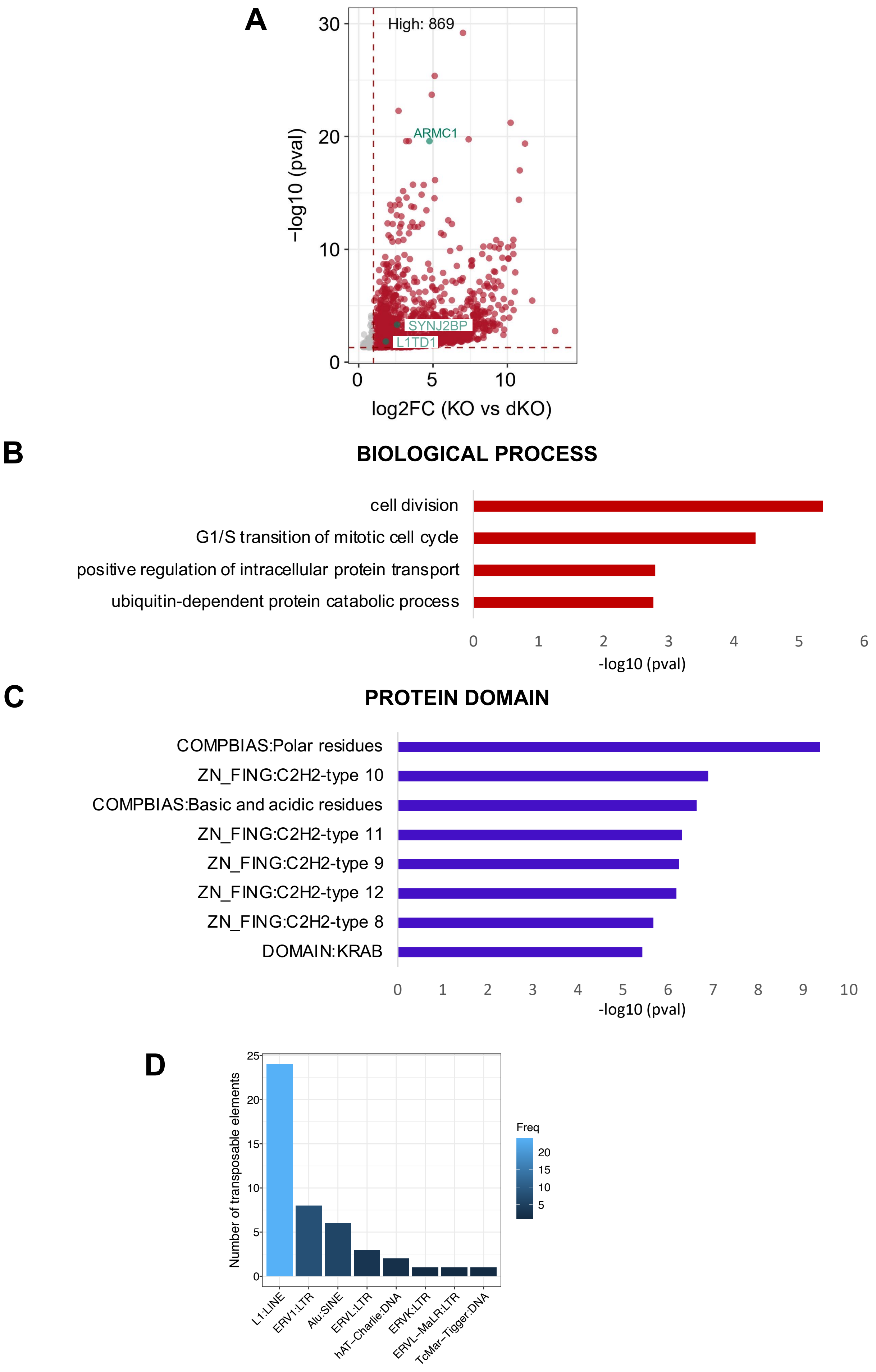

### Supplemental Figure S4

#### Supplementary Figure S4

A

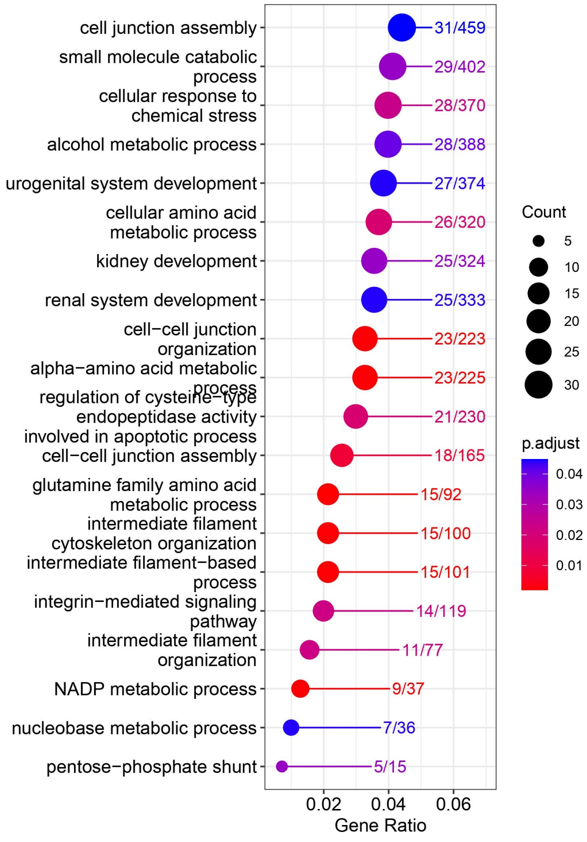


B

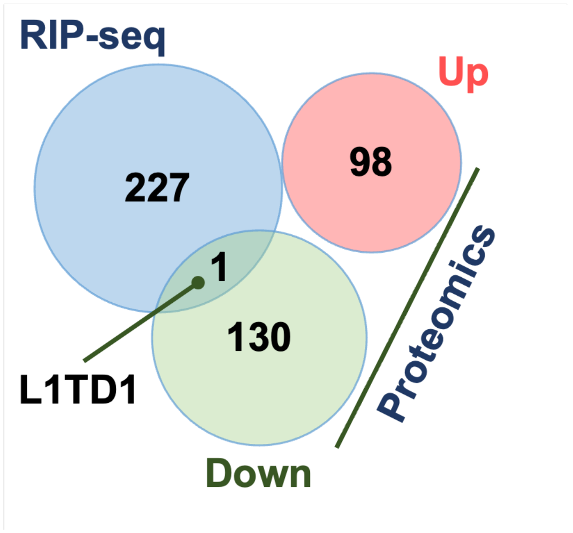

### Supplemental Figure S5

#### Supplementary Figure S5

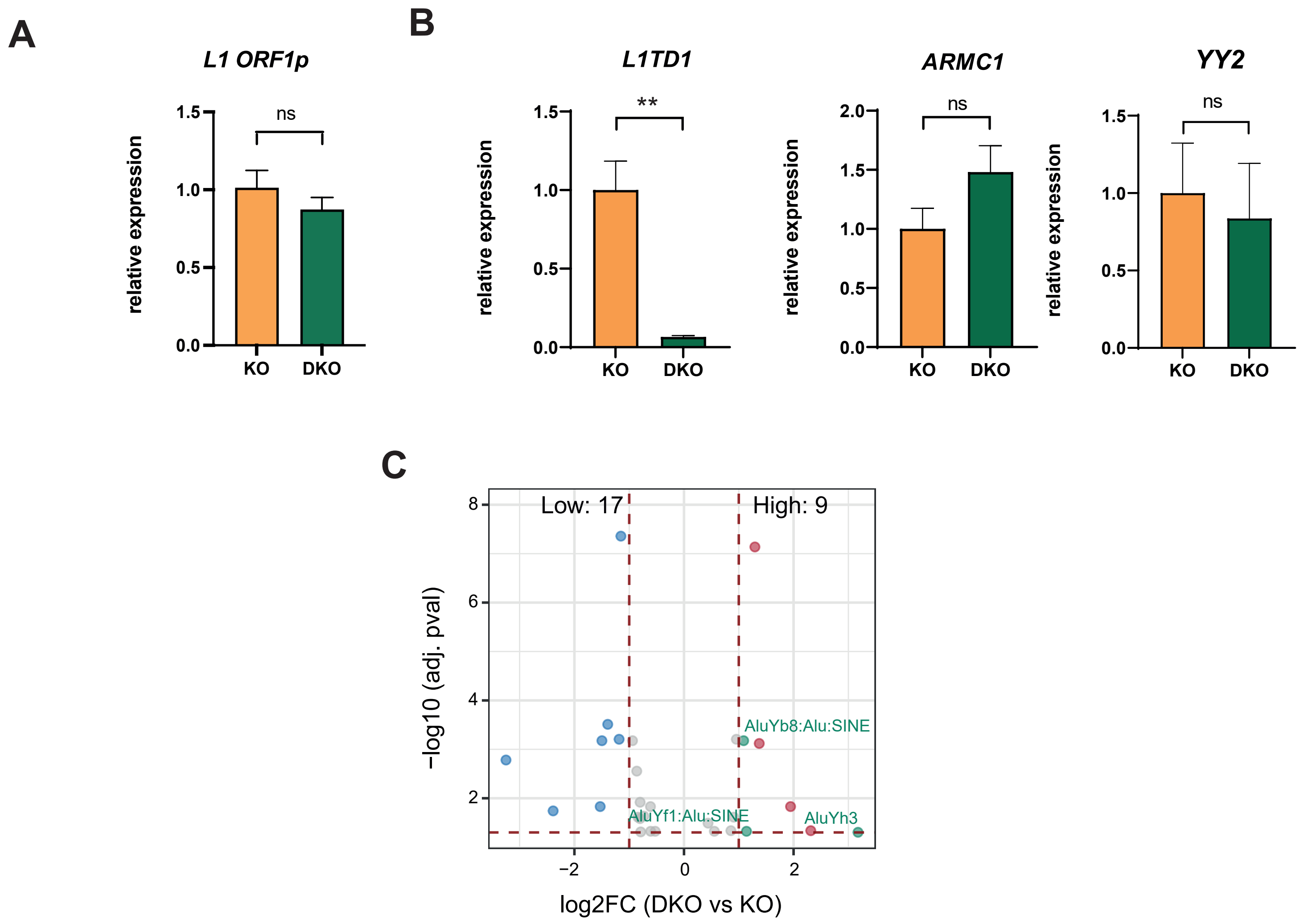

### Supplemental Figure S6

##### Supplementary Figure S6

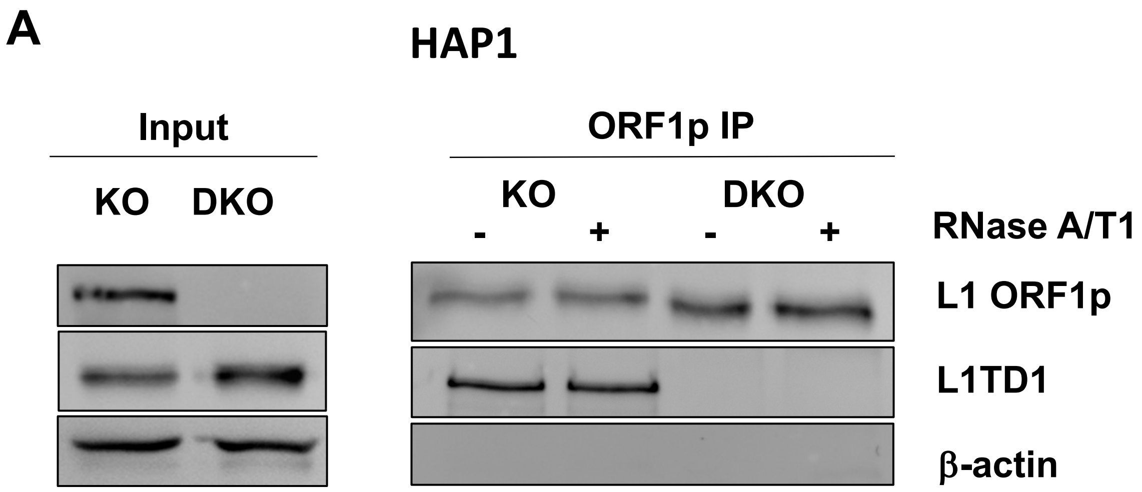


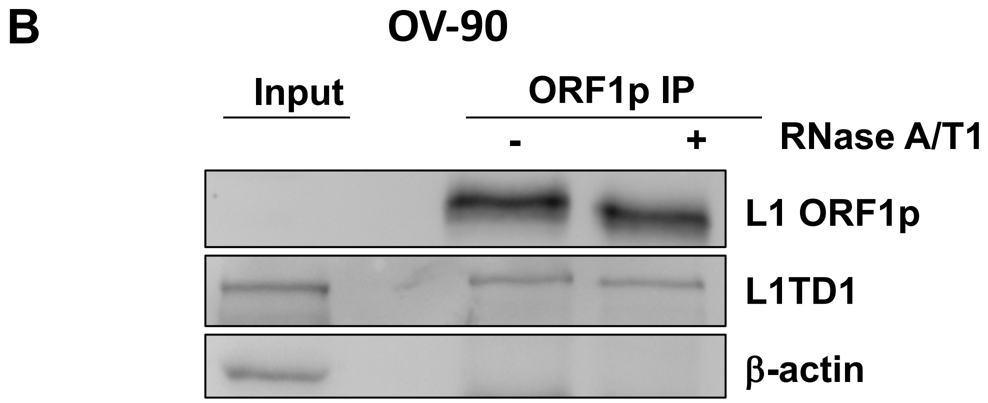


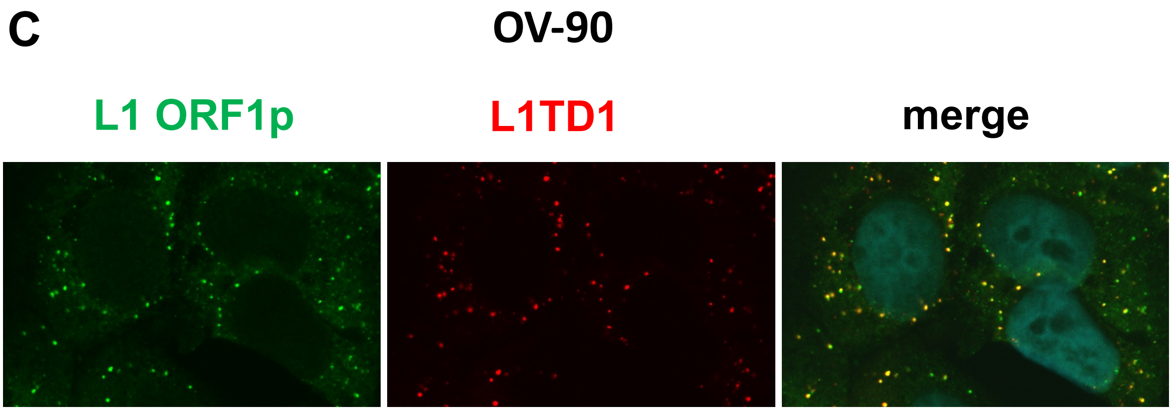

### Supplemental Figure S7

#### Supplementary Figure S7

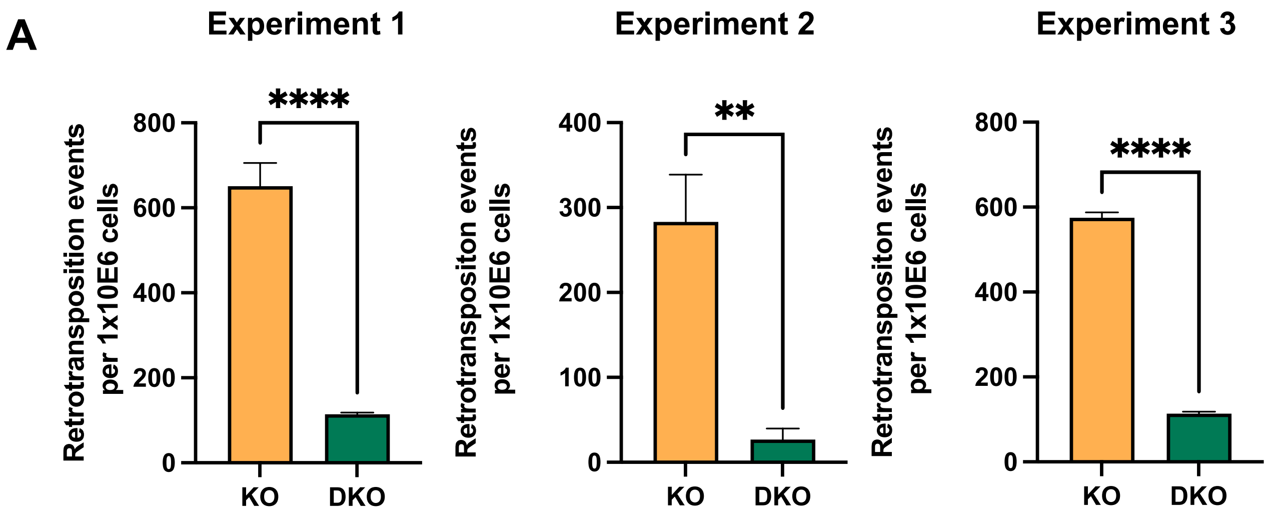


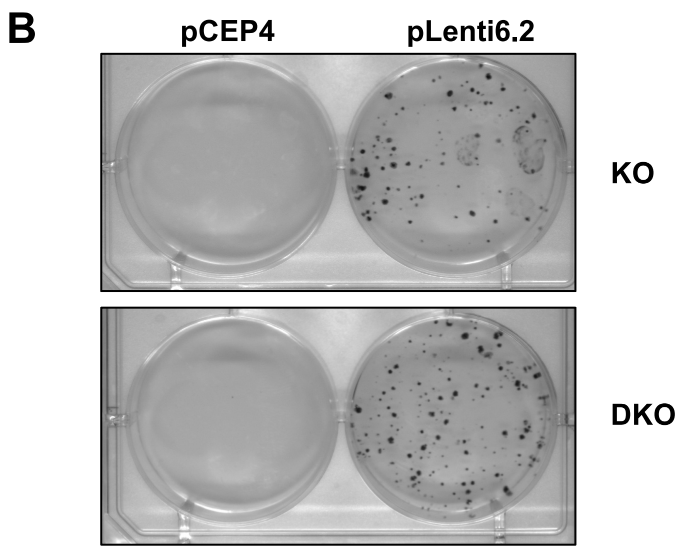


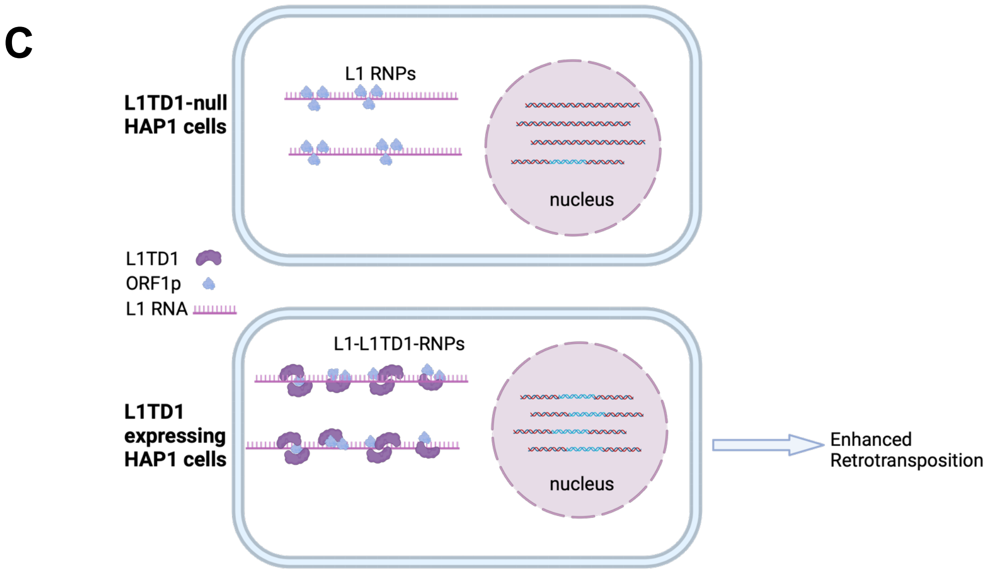
